# Abiotic niche predictors of long-term trends in body mass and survival of Eastern Himalayan birds

**DOI:** 10.1101/2022.08.25.505219

**Authors:** Akshay Bharadwaj, Ritobroto Chanda, Aman Biswakarma, Bharath Tamang, Binod Munda, Dambar K Pradhan, Mangal K Rai, Shambu Rai, Umesh Srinivasan

## Abstract

The synergistic impacts of climate change and habitat degradation threaten tropical species worldwide. However, how species’ abiotic niches affect their demographic vital rates and phenotypic changes under anthropogenic change remains poorly understood. Using an 11-year mark-recapture dataset from primary and selectively logged forest in the Eastern Himalayas, we investigated how temperature-humidity niche characteristics predicted body mass and survival trends in understorey insectivorous birds over time in each habitat. Our results show that logged forest is hotter and drier than primary forest, and the arthropod community shows dramatic shifts in composition upon selective logging. In understorey insectivores, the degree of dissimilarity between species-specific primary and logged forest niches was strongly and negatively correlated with survival and body mass trends in logged forest. Here, we show that temperature-humidity niche shifts in response to anthropogenic habitat modification can impact demographic vital rates and body condition crucial for population persistence. This work has the potential to inform prompt, targeted conservation efforts toward species that are the most threatened in a warmer and more degraded world.

## Introduction

Climate change and habitat degradation are the two greatest threats to tropical biodiversity today (Pimm, 2008; Pringle et al., 2012). Not only are global temperatures rising rapidly due to climate change, the anthropogenic modification of tropical forest (e.g., by selective logging and conversion to agriculture) further compounds climate impacts; modified habitats reach far higher temperatures than natural forests (Jucker et al., 2020) and lose the buffering effects of the forest canopy (De Frenne et al., 2019). There is also a general consensus on the expected effects of climate change and forest modification on the structure and function of ecological communities — large-bodied, *k*-selected species and those at higher trophic levels decline in abundance with selective logging and habitat fragmentation (Burivalova et al., 2015; Duffy, 2003; Gray et al., 2007; Hamer et al., 2015; Petchey et al., 1999).

While the impacts of these two factors on biodiversity have received great attention, these impacts have been mostly considered separately, notwithstanding the fact that species are simultaneously exposed to, and therefore respond to, both climate change and habitat degradation concurrently (Srinivasan & Wilcove, 2021). Further, most research on the impacts of climate change and forest loss often examines biodiversity responses in terms of range shifts (Freeman et al., 2018), changes in the population sizes of species (Kim et al., 2022; Şekercioğlu et al., 2012), and the structure of ecological communities (Newmark, 2006). Little is known about the compounding effects of climate change and habitat modification on demographic vital rates such as survival, and phenotypic characteristics, such as body mass (Cosset et al., 2019; Messina et al., 2021; Srinivasan, 2019). Such an understanding, however, remains crucial because: (a) vital rates determine the long-term persistence and viability of populations; (b) studying phenotypic changes over time can help us understand the underlying drivers behind shifts in community structure and vital rates; and (c) quantification of an objective, reliable estimate of a species’ vulnerability to extinction — using niche parameters—enables effective use of resources to best conserve biodiversity.

All species occupy distinct ecological niches, defined as the sum of the habitat requirements, environmental conditions and behaviours that enable a species to persist at a location (Jirinec et al., 2021; Ruegg et al., 2021). All else being equal, species with large fundamental abiotic niches, quantified as a function of environmental parameters such as temperature and humidity, should be more adaptable to habitat modifications that alter the abiotic environment (Moritz & Agudo, 2013). In contrast, niche specialists with narrow temperature-humidity niches are likely to be more sensitive to environmental change (Clavel et al., 2011). Temperature and humidity have especially been shown to determine species, and even individual, responses to habitat change and the resulting alteration in climatic variables (Bay et al., 2021; Frishkoff et al., 2016; Kim et al., 2022; Neate-Clegg et al., 2021). For instance, outright habitat loss from conversion to agriculture leads to an average 7.6°C increase in temperature and a reduction in humidity relative to primary forest (Fick & Hijmans, 2017). Even relatively minor forest degradation (for instance, through selective logging) raised mean temperatures by 2°C and maximum temperatures by 5.8°C (Srinivasan & Wilcove 2021).

Selective logging is the predominant logging regime in the tropics (Edwards & Laurance, 2013), and results in logged forest occuring in a continuum of varying tree densities throughout the world (Burivalova et al., 2015). Although not leading to outright habitat loss, selective logging leads to habitat degradation and the associated impacts on plant and animal communities (Halaj et al., 2008; Padmawathe et al., 2004; Velho et al., 2012). Logging can directly affect species; for instance via the felling of trees leading to reduced availability of nest spaces and refugia and increased predation risks (Tuff et al., 2016). Logging can also impact species indirectly through changes in microclimate, an important determinant of species presence across plant and animal taxa (Boyle et al., 2021; Zellweger et al., 2020). Microclimatic characteristics and diversity are in turn linked to the physical structure of the tree community (Han et al., 2022) and consequently, sensitive to selective logging (Aleixo, 1999; Thiollay, 1992).

Altered thermal environments can also affect phenotypes. A morphological change that has been postulated to occur under climate warming is the reduction of body size (Watt et al., 2010). This follows from Bergmann’s rule, based on the fact that a higher surface area:volume ratio translates to better heat loss, leading to the expectation that warming climates should favour smaller body sizes (Buskirk et al., 2010; Gardner et al., 2011). The evidence, however, shows mixed support for this ‘rule’ with some species showing reductions in body size, while others showed increased sizes with rising temperatures (Jirinec et al., 2021; Teplitsky & Millien, 2014). This may be because the adaptability of species to climate change and the determinants of this adaptability are yet to be fully understood. An understanding of the drivers of adaptability of a species to climate change and habitat degradation is, therefore, crucial to forecast the vulnerability of various species present in an ecosystem. Furthermore, the ability of a species to adapt to altered abiotic conditions via reduction in body size is likely to also be linked to long-term survival in modified habitats (Møller & Szép, 2002; Rioux Paquette et al., 2014).

We studied year-round resident bird species in the Eastern Himalaya Global Biodiversity Hotspot for an 11-year period, using mark-recapture in both primary and logged forest plots. The Eastern Himalayas is among the world’s most biodiverse regions (Grenyer et al., 2006; Pandit et al., 2014), and presently faces the twin threats of selective logging and climatic warming. Furthermore, Eastern Himalayan bird species, especially the understorey insectivores, are adapted to narrow thermal ranges and microhabitats, potentially making them extremely vulnerable to anthropogenic change (Jetz et al., 2007; Powell et al., 2015; Srinivasan et al., 2018).

In this study, we asked:

a. Are there abiotic and biotic differences between logged and primary forest environments? **We expected logged forest to be hotter and drier than primary forest habitats. As a consequence, we also expected shifts in the biotic communities of arthropods in selectively logged forests, due to their thermally-sensitive and ecothermal nature (Renault et al., 2022).**
b. Does the degree of dissimilarity between species’ primary and logged forest niches correlate with their body mass and/or survival trends over time in logged forest? **We expected that the more dissimilar the primary and logged forest abiotic niche of a species is, the steeper the body mass and survival declines of that species in logged forest. In other words, we expect to see a positive correlation between the degree of overlap between primary and logged forest niches, and species’ body mass and survival trends over time in logged forest.**

## Methods

### Study area and field sampling

We sampled primary and logged montane broadleaf wet evergreen forest at 2000m ASL in Eaglenest Wildlife Sanctuary, West Kameng district, Arunachal Pradesh, India. Six forest patches were selected for sampling based on interviews with persons involved in logging operations (three each in primary and logged forest; a total of 9ha in each habitat; plot centroids were separated by an average of 721 ∓ 301m SD; Fig S1). Variation in tree densities on these plots tallied exactly with semi-quantitative estimates of timber extraction (Srinivasan, 2013). On average, tree densities in primary forest plots (∼200 trees per ha) were more than double those in logged forest (∼75 trees per ha).

In each plot, a team operated 24-48 mist nets (12m length, 4 shelf, 16mm mesh size; 158 nets in total) from 0500 to 1200h for three consecutive days during the early breeding season (April-May) each year, from 2011-2021. Sampling was not possible in 2020 because of a nationwide lockdown caused by the COVID-19 pandemic. Mist-netting was conducted in three cycles, each spanning three days each in every plot. Nets were set up systematically at the same locations each year within well-defined study plots, with neighbouring nets placed ∼40m apart (Fig S1). No mist nets were placed along roads or trails, but a narrow line was cleared in the understorey prior to the deployment of mist-nets. Because sampling was done during the early breeding season, all captured small passerine birds were adults and were found in small, distinct territories very little movement between plots (Srinivasan & Wilcove, 2021). All captured individual birds were ringed with a uniquely numbered aluminium ring, weighed and then released.

### Differences in overall temperature-humidity profiles in primary and logged forest

To measure the temperature and humidity associated with each capture of an individual bird, a single DS 1923 iButton Hygrochron (Maxim Integrated) temperature-humidity logger was placed at the location of each mist net (158 logger locations in total). Temperature-humidity loggers were programmed to record both temperature and relative humidity at 10-minute intervals. Loggers were deployed along with the mist nets throughout the mist-netting conducted in 2021, which was conducted in three cycles spanning three days each in every plot. We returned to mist net in each plot roughly after every 15 days, a timeframe in which there are no major changes in climate in the relatively aseasonal north-eastern part of India (Srinivasan et al., 2018). Because the order of deployment in the plots was randomised, we do not expect any biases in microclimate characterisation across habitats. A previous behavioural study which also measured microclimates from the same plots in an earlier period (March-April) in 2021 showed qualitatively the same temperature-humidity differences between primary and logged forest (Aggarwal et al., 2023).

Each plot had 25-28 mist nets, and correspondingly, 25-28 temperature humidity loggers (one deployed at each mist net). Each bird capture was then associated with the closest temperature and humidity reading in time from the mist net of capture. The bird species we select for our analyses are insectivorous birds found in the understorey/midstorey (where our temperature-humidity loggers are deployed). They do not move up vertically into higher forest strata such as the canopy, thereby allowing an accurate estimation of their abiotic niche use. Unlike capture efforts (which span 2011 to 2021), temperature-humidity data associated with bird captures in primary and logged forest was only measured in 2021.

To analyse the temperature and humidity profiles of primary and logged forest, we used a mixed effects model in the lme4 package (Bates et al., 2015) in R (R. Development Core Team, 2013) parameterised as follows:

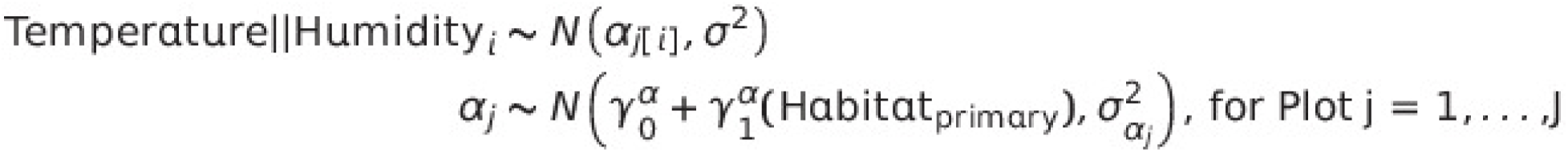

where habitat was the fixed variable with two levels (primary and logged) and plot was a random variable with six levels (three plots each in primary and logged forest). From these models, we report the differences in temperature and humidities between primary and logged forest (i.e., the effect size) and 95% confidence intervals.

### Differences in arthropod community composition between primary and logged forest

Like all ectotherms, arthropods are sensitive to the temperature and humidity changes (Burraco et al., 2020) and make up the diets of all our study species (Supriya et al., 2020). Therefore, to better understand biotic changes accompanying selective logging, we studied the arthropod community compositions in primary and logged forest.

We used three arthropod sampling techniques at 25-28 points within each plot to sample the arthropod community in primary and logged forest (55 sampling stations in primary forest and 53 sampling stations in logged forest). We accounted for ground-dwelling, flying and understorey foliage arthropods through the use of sticky traps, pitfall traps and branch-beats.

For ground-dwelling arthropods, we placed pitfall traps containing detergent water for 48 hours at each sampling station. For flying arthropods, we used sticky traps - a cardboard rectangle 150 x 200mm with an adhesive coating on both sides. These non-toxic sticky traps contained an aromatic substance to attract insects, and were hung approximately 1.5m from the ground for 48 hours at each location. Lastly, we used branch beating for sampling foliage arthropods. For branch beating, we stitched a white linen cloth in the shape of a funnel around a metal ring 1m in diameter at the larger end of the funnel and with a container attached to the thinner end of the funnel. At each station, we placed a randomly selected branch within this funnel, beat it ten times with a standard-sized stick and sprayed commercially available insecticide spray. This momentarily paralyzed the arthropods, allowing us to collect them in the container affixed to the narrow end of the funnel.

All collected arthropod samples were photographed with scale using a Nikon P900 camera. Each arthropod within the sample was identified to the order level.

### Selection of bird species for analysis

We selected species suitable for analyses based on the following criteria:

1. Selected species were understorey insectivores that breed on our sampling plots (migrants were excluded).
2. Selected species had sufficient recapture rates to provide robust estimates of species survival in both primary and logged forest.
3. Selected species had sufficient temperature-humidity data associated with their captures to allow the modelling of the abiotic niches.

The list of all selected species is reported in Table S1.

### Temperature-Humidity niche analysis

To estimate the temperature-humidity niches of different species we used the *adeHabitat* package (Calenge, 2006) in Program R (Core Team 2022), to model the temperature-humidity niche using a Utilisation Distribution (UD) kernel approach (Calenge, 2006; Fig 1). While this method was initially developed to analyse the utilisation of various parts of the home ranges of radio-tracked animals using a Kernel Density Estimate (KDE), we use it to model the use of the temperature-humidity space for each species, in primary and logged forest (Fig 1). Through this approach, for each species, we generated separate temperature-humidity KDEs for primary and logged forest separately, and subsequently calculated the Utilisation Density Overlap Index (UDOI) — a standard measure of UD overlap based on the Hurlbert’s index of niche overlap (Hurlbert, 1978; Fig 1). UDOI (henceforth ‘niche overlap’) is an individual covariate associated with the capture history of each individual, and is calculated separately for each species. All individuals of a particular species are therefore assigned the same UDOI value in the mark-recapture models (see Srinivasan et al., 2015).

**Figure 1:**
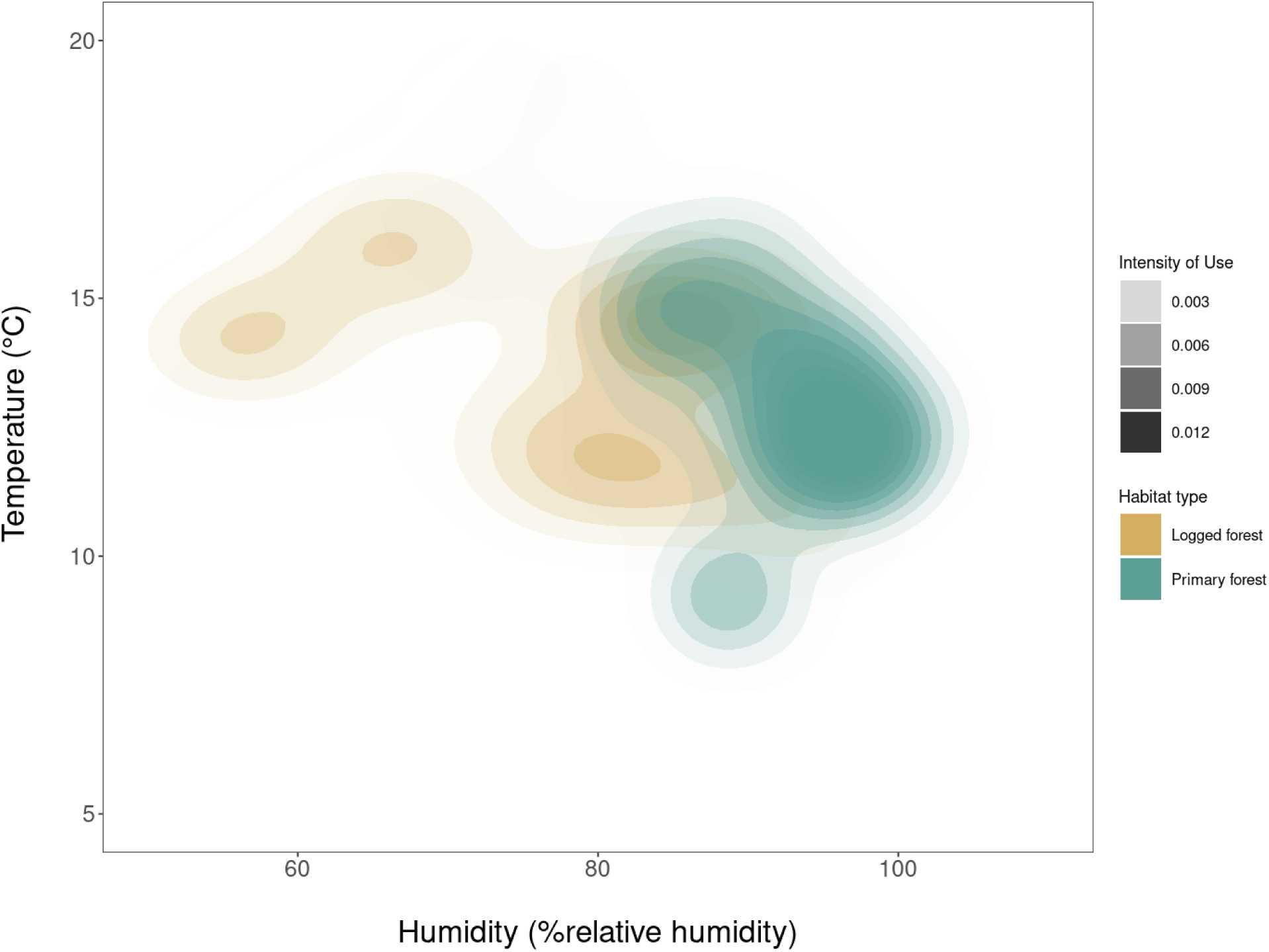
**Example of the temperature-humidity niches of Yellow-throated Fulvetta (*Pseudominla cinerea*) in primary and logged forest, visualised as a Utilisation Distribution kernel. The abiotic niche for this species is generally more moist and slightly cooler in primary than in logged forest. The intensity of use of a given region in temperature-humidity space is indicated with a gradient of shading, opaque representing region of most use and transparent representing regions of no use.**

### Body mass analysis

First, we standardised all body masses to a species-wise z-score to account for the fact that species which are larger can potentially lose more absolute body mass than species which are inherently small. We then used a generalised linear model to examine the relationship between body mass, time and niche overlap.

We used generalised linear models to examine how the average body mass of birds of each species changed over time in primary and in logged forest. This was modelled as follows:

**Z-score ∼ Year * Habitat * Niche_Overlap**, where

1. Z-score = Z-score of the body mass (continuous variable)
2. Year = Year of study (continuous variable)
3. Habitat = habitat type (categorical variable with two levels; primary and logged)
4. Niche_Overlap = Niche overlap between primary and logged forest niches (continuous variable)

To visualise our results we created a heatmap whose axes was time and niche overlap, and the color of the grid cells indicated the Z-score predicted by our model. Niche overlap, which is a continuous variable, was split into 10 different equally-spaced values within the naturally occurring range of values.

To visualise the difference in body mass trends between species with highly overlapping and highly disjunct primary and logged forest niches, we plotted the predicted Z-score over time for these species only.

### Survival analysis

We estimated the apparent survival (phi) for each species in each type of habitat using the Cormack-Jolly Seber (CJS) mark-recapture model (Cormack, 1964; Jolly, 1965; Seber, 1965), to estimate adult apparent survival in primary and logged forests separately (Fig S1). Apparent survival is an amalgam of true survivorship and site fidelity (i.e., death and emigration are indistinguishable; Barbour et al., 2013). Therefore, an annual apparent survival estimate of 0.60 indicates a 60% probability of an individual in the population surviving *and* remaining in the same location from one year to the next. For each species, to analyse how apparent survival changed over time, we first created a capture history for each individual bird spanning eleven years. Multiple captures of the same individual within a single year (e.g., on consecutive sampling days) were not considered recaptures (i.e., all captures within a single sampling season were collapsed into a single event in the capture histories). We then pooled capture histories from the three primary forest plots to represent populations sampled in primary forest and pooled capture histories from all logged plots to represent populations sampled in logged forest. We pooled the data to estimate apparent survival in primary and logged forests for each species because plot-level data was insufficient to adequately and reliably estimate survival for each species in each plot.

The CJS model is an open-population model that estimates apparent survival while accounting for imperfect detection (i.e., capture probability < 1). Using the *R2ucare* package (Gimenez et al., 2018) in Program R (Core Team, 2013), we first ran tests to ensure that the CJS models fitted the capture history data well. We ran two models separately — one in which recapture probability was constant throughout the study period (across all sampling occasions), and another in which recapture probability was allowed to vary with sampling occasion. In both models, apparent survival was constrained to vary unidirectionally over time (but the slope of survival over time was allowed to be zero). We chose the best model (between time-variant and time-invariant recapture probabilities) by using the small sample size-corrected Akaike’s Information Criterion (AICc). The best-performing model was used, and it was as follows:

**Phi(∼Year * Habitat * Niche_Overlap)p(∼1)**, where

1. Phi() = apparent survival (continuous variable between 0 and 1)
2. Year = Year of study (continuous variable)
3. Habitat = habitat type (categorical variable with two levels; primary and logged)
4. Niche_overlap = niche overlap between primary and logged forest niches (continuous variable)
5. p(∼1) = capture probability (remains constant in the best performing model)

Using the estimates from this model, we created a heatmap whose axes was time and niche overlap, and the color of the grid cells indicated the survival predicted by our model. Niche overlap, which is a continuous variable, was split into 10 different equally-spaced values within the naturally occurring range of values.

To visualise the difference in body mass trends between species with highly overlapping and highly disjunct primary and logged forest niches, we plotted the predicted survival over time for these species only.

## Results

### Temperature and humidity in primary and logged forest

Over the entire 24-hr daily period, primary forest was -0.87 degC cooler than logged forest plots (95% CI = [-2.17, 1.97]; marginal R2 = 0.0001; conditional R2 = 0.10). However, while the mean temperature does not differ significantly between the two, it hides differences in the daily temperature regimes in primary and logged forest. Logged forest was more variable than primary forest with lower temperatures at night, warming more rapidly during the day, up to 4.4℃ warmer than primary forest at mid-day (mean difference in temperatures between primary and logged forest at 1300hrs = 0.83℃; Fig 2a).

**Figure 2:**
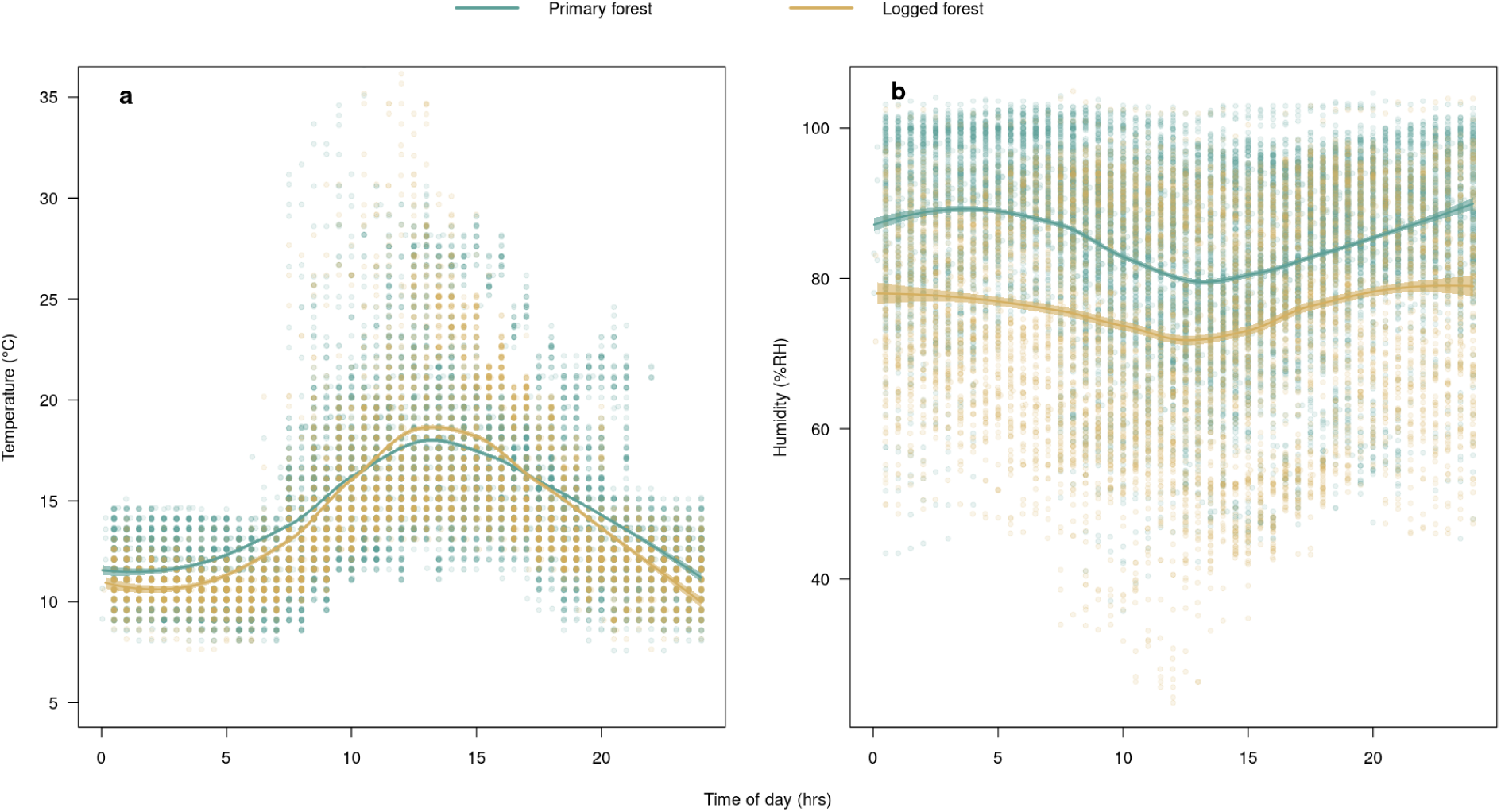
(a) Daily temperature (in. ℃**) and (b) daily relative humidity (%) trends in primary (green) and logged forest (brown) over a twenty-four hour period. Logged forest has, on average, more variable temperatures and is consistentely and considerably drier than primary forest. Solid lines indicate trendlines from locally estimated scatterplot smoothening (LOESS), and the polygons around the lines represent the 95% confidence interval around the trendline.**

Regardless of the time of the day, on average, logged forest was less humid than primary forest (difference in average humidity between primary and logged forest = 8.73%; 95% CI = [-3.73, 21.20]; marginal R^2^ = 0.08; conditional R^2^ = 0.39; Fig 2b).

### Arthropod Community Composition

Primary and logged forest differed significantly in athropod community composition (NMDS, MoristaHorn index, stress = 0.20; perMANOVA: *F*_1,106_ = 24.69, *p* < 0.01, R^2^ = 0.82). In general, primary forests harboured more arachnids, lepidopterans and hemipterans while logged forest insect communities had more flies (Fig 3). The total abundance of arthropods did not differ significantly between primary and logged forest (Poisson ANOVA, *z*_1,106_ = 1.77; *p* = 0.08), although there were trap-specific differences observed. Foliage arthropod density was higher in primary forest (branch beats; ANOVA, *t*_1,106_ = 4.22; *p* < 0.01) while density of arthropods in flight was higher in logged forest (sticky traps; Z_1,106_ = -5.12, p < 0.01; Aggarwal et al., 2023). No difference was observed between terrestrial arthopod abundances in primary and logged forest (pitfall traps; Z_1,106_ = 0.33, p = 0.75; Aggarwal et al., 2023).

**Figure 3:**
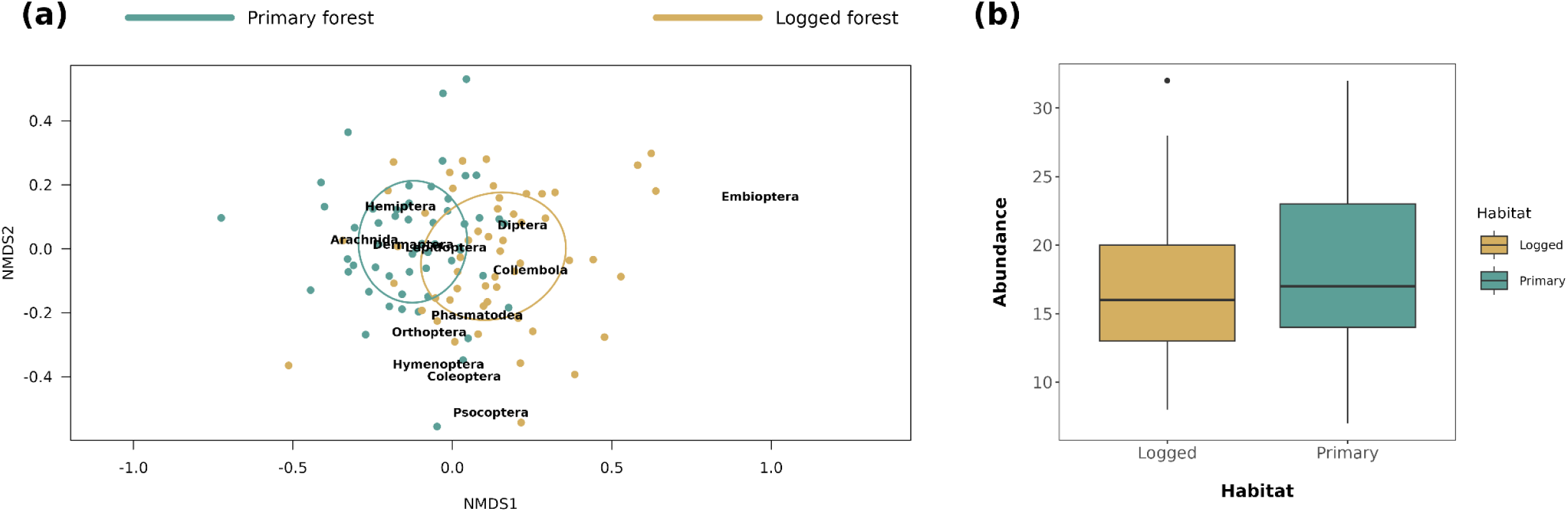
**(a) An NMDS visualisation of the arthropod community composition in primary and logged forests at our study site. Primary forest plots had more arachnids, lepidopterans and hemipterans while logged forest insect arthropod communities have a greater proportion of dipterans; (b) The abundance of arthropods in both logged and primary forest are not significantly different (Poisson ANOVA, *z*_1,106_ = 1.77; *p* = 0.08). However, there are trap-wise differences between primary and logged forest arthropod communities, corresponding to the higher abundance of arachnids and dipterans in primary and logged forest, respectively (Aggarwal et al., 2023).**

### Preliminary mist-netting results

During the 11-year study period, we recorded 9,316 captures of 6,340 individuals of 135 species. Based on three selection criteria (breeding, enough data for capture-recapture analysis and enough data for temperature-humidity niche modelling) we selected 17 species for further analyses. In total, we recorded 5,428 captures of 3,519 individuals belonging to these 17 species.

### Body mass trends

Our analysis suggests a significant effect of climate and land-use change on the body mass trends of birds over time. In logged forest, body mass trends over time showed a positive relationship with niche overlap (*β*_time*niche_overlap_ = 0.03, 95%CI = [-0.01, 0.08]; *β*_time*niche_overlap*habitat_primary_ = -0.04, 95%CI = [-1.10, 0.02]; Fig 4a-b). Species with greater niche dissimilarities between their primary and logged forest niches also had steeper survival declines over time in a logged forest (Fig 4b). The model parameters for the estimates and confidence intervals are reported in Table S2.

**Figure 4:**
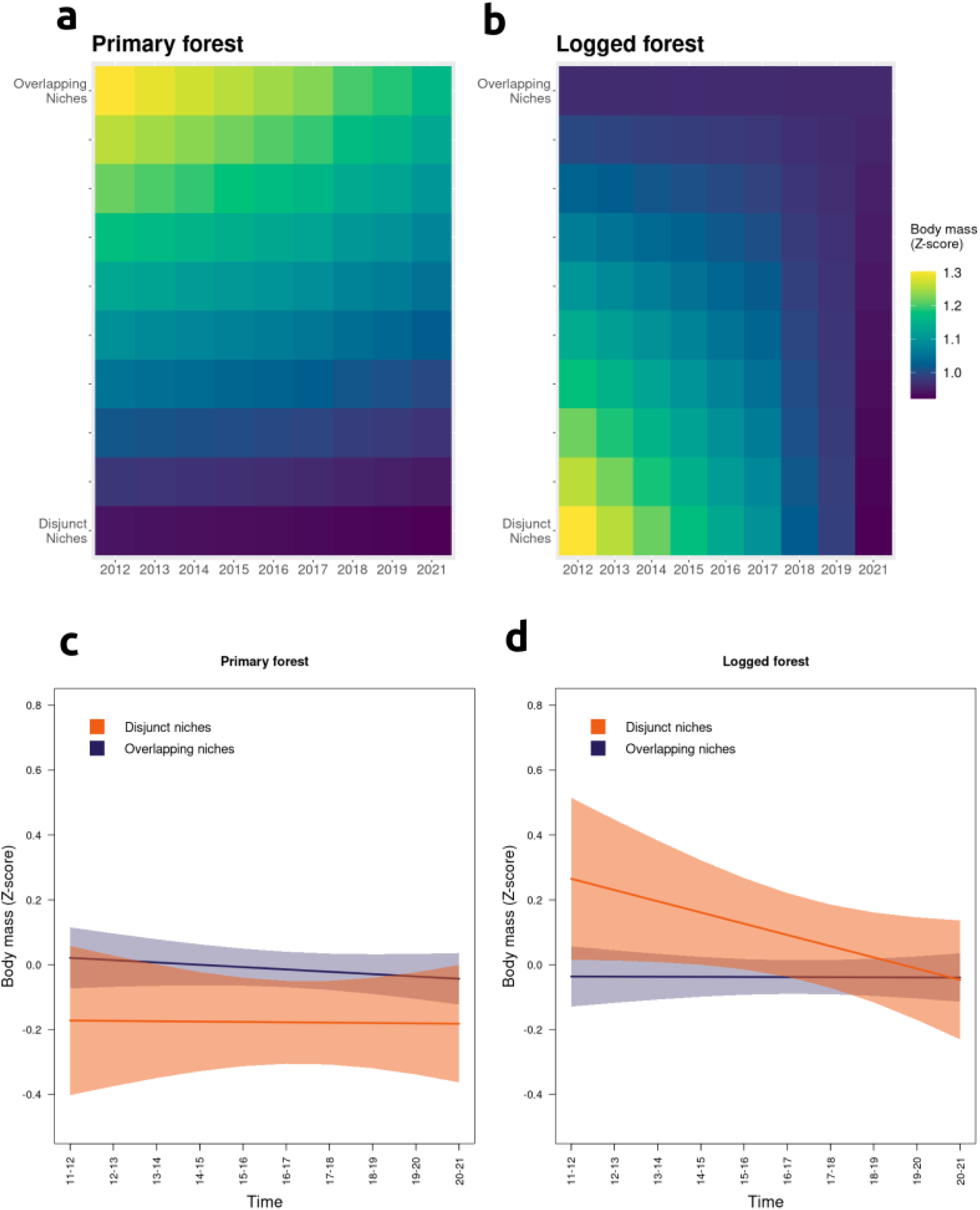
**(*Top row*) A heat map showing the relationship between body mass, time and niche overlap in (a) primary and (b) logged forest. Cells with darker hues of blue indicate lower survival, while those with hues of green have higher survival; (*bottom row*) The relationship between survival and time for species with high (blue) and low (orange) niche overlap between their primary forest and logged forest niches.**

### Survival analysis

Individuals that were captured on more than one habitat type (primary or logged) were excluded from further analyses (3.9% of all captured individuals). Models with constant recapture probability performed better for all species, in comparison to one with time-varying capture probability. The model parameters for the estimates and confidence intervals are reported in Table S3.

Niche dissimilarity (i.e., the degree to which a species’ temperature-humidity niche changes with logging) is strongly related to survival trends in logged forest (*β*_time*niche_overlap_ = 0.23, 95%CI = [0.001, 0.47]; *β*_time*niche_overlap*habitat_primary_ = -0.37, 95%CI = [-0.70, -0.03]; Fig 5a-b). Species with greater niche dissimilarities between their primary and logged forest niches also had steeper survival declines over time in a logged forest (Fig 5b).

**Figure 5:**
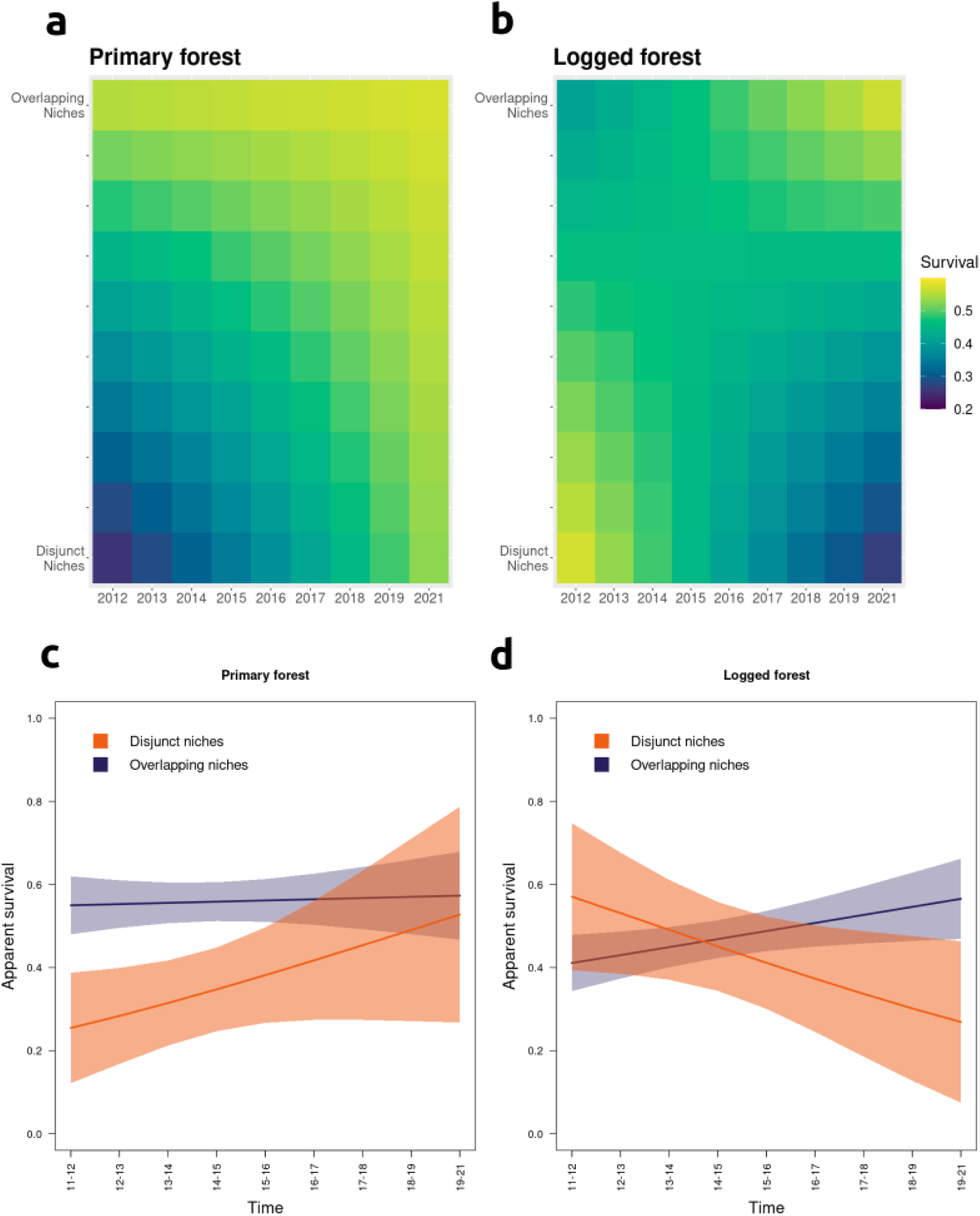
**(*Top row*) A heat map showing the relationship between survival, time and niche overlap in (a) primary and (b) logged forest. Cells with darker hues of blue indicate lower survival, while those with hues of green have higher survival; (*bottom row*) The relationship between survival and time for species with high (blue) and low (orange) niche overlap between their primary forest and logged forest niches.**

## Discussion

We examined differences in the abiotic and biotic environments for 17 species of understory insectivorous birds in primary and selectively logged forest in the Eastern Himalaya. We analysed how the temperature-humidity niche sizes of understorey bird species were related to body mass and survival trends over an 11-year period in primary and logged forest. Primary and logged forest had different temperature-humidity profiles and differed in the composition of arthropod communities. Importantly, less than four percent of all captured birds were captured in two different plots, indicative of their small territory sizes and site fidelity during the breeding season, irrespective of the habitat type that the individual is found in (Ibarra-Macias et al., 2011; Srinivasan & Wilcove, 2021).

As hypothesized, we found a strong positive relationship between the degree of niche dissimilarity (i.e., the degree to which a species’ temperature-humidity niche changes with logging) and survival rates and body mass trends over time. In other words, species with very different temperature-humidity niches in primary and logged forest experienced survival and body mass declines in logged forest, while those with high degree of niche overlap between primary and logged forest have near-constant or increasing survival and body mass over time (Fig 4, Fig 5), even after logging.

Body size is an important and widely-studied life-history trait that greatly influences thermoregulation by individuals/species (James, 1970). Habitat-specific differences in a species’ body size (specifically, body mass) can be the result of two possibly concurrent processes: (a) the consistent reduction in body mass of each individual of the species and/or (b) the accumulation of lower quality (in this case, lighter) individuals in logged forest.

Heat dissipation theory postulates that tropical animals are limited energetically by the rate at which body heat is lost (Speakman & Król, 2010), and purely based on thermodynamic considerations, smaller body sizes should be selected for in progressively warmer conditions (Prokosch et al., 2019). The Eastern Himalaya has a relatively aseasonal climate which selects for thermal niche specialist species (Srinivasan et al., 2018). Not only is the Eastern Himalaya warming three times as fast as the global average (Shrestha et al., 2012), but forest degradation is concomitantly removing the buffering effects of the primary forest canopy, making logged forests more thermally variable, and significantly hotter and drier than primary forest (De Frenne et al., 2019; Figs 2a-b). Indeed, our *a priori* expectation was that thermally-stressed specialist species would undergo steeper declines in body mass over time, in keeping with the thermoregulatory capacity hypothesis. In other words, species with more different primary and logged forest niches would undergo steeper declines in body mass over the study period. Indeed, this is what we observe - body mass over time is negatively correlated with niche overlap (Fig 4a).

Several bird species elsewhere are indeed undergoing other morphological changes over time, likely because of climate-driven thermal constraints (Jirinec et al., 2021; Messina et al., 2021). Specifically, birds can undergo changes in the morphometry of organs and appendages that serve thermoregulatory (Tattersall et al., 2017) and locomotory functions (Weeks et al., 2020), without drastically changing overall body mass. In the rainforests of the Amazon, for example, climate change appears to have selected for progressively longer wings in comparison to body mass, which suggests that birds are adapting toward relatively longer appendages to offset thermal stress (Jirinec et al., 2021). Therefore, in addition to progressively lighter body mass in logged forests, birds within primary and logged forests would likely exhibit differences in their mass:wing ratio, a hypothesis that remains to be tested.

In addition to thermal stressors, resource availability also plays a role in determining body size (McNab, 2010). Since we were interested in insectivorous understorey birds, we also measured biotic factors determining body sizes such as arthropod composition and abundance (Cox et al., 2019). Arthropod community composition showed a considerable shift with selective logging. This result highlights that a possible confounding variable hitherto unexplored at our study site is the dietary niche breadth of birds (Bond & Lavers, 2014; Hansen et al., 2019). An assessment of dietary niche space (i.e., its size and shift in response to selective logging) would help in better understanding the various mechanisms underlying body mass declines with logging (Edwards et al., 2013; Hamer et al., 2015; Messina et al., 2021). Furthermore, we investigate only the abiotic changes brought about by selective logging. Investigating the other impacts of logging, such as an altered availability of nest spaces and predator-prey interactions will shed more light on the mechanisms impacting species over time (Tuff et al., 2016).

We also acknowledge that the trends we observe in body mass declines over time in the logged forest can occur through the movement of lighter, “low-quality” individuals to the logged forest plots (Calsbeek & Sinervo, 2002). Such a mechanism, arising from a despotic distribution, would be expected to result in a situation where the body masses of individuals (within a species) in logged forest would, on average, be lower than for individuals in primary forest (Andren, 1990). In the absence of long-term processes such as climate change, body mass differences between primary and logged forest would be expected to be stable. However, we report consistent long-term declines in body mass of in logged forest of species that show little niche overlap between the two habitats, likely as a result of warming-induced morhphological change (Jirinec et al., 2021).

The characteristics of a species’ niche determines how it interacts with the larger ecosystem and shapes its individual and population-level responses to ecological disturbances (Bellwood et al., 2019; Neate-Clegg et al., 2021; Voigt et al., 2007). Previous studies have suggested that species with larger niches (i.e., generalists) can inhabit a greater range of environments by drawing from a wider pool of resources. This makes them more resilient to climate change and less likely to face population declines in comparison to species with smaller niches (Clavel et al., 2011; or, specialists; Swihart et al., 2003). Specialist species use a narrow range of resources and/or habitats, and changes in the availability of both can have drastic effects on their populations since they are unable to exploit alternative habitats or resources (Graham et al., 2011).

Studies investigating the effect of niche characterstics on sensitivity to climate change have thus far used occupancy data and theoretical models to estimate niche sizes (Loiselle et al., 2010; Rinnan & Lawler, 2019). The direct impact of climatic niche parameters on the long-term survival of species has seldom been tested in the field. Previous studies on tropical rainforest species show that vulnerability is strongly predicted by ecological specialisation (Sekercioglu, 2011), and that understory insectivores are in particular peril due to anthropogenic change because of their limited dispersal ability (Moore et al., 2008; Powell et al., 2015) and sensitivity to the opening up of the canopy (Patten & Smith-Patten, 2012; Pollock et al., 2015).

We found that the degree of niche dissimilarity between primary and logged forests was strongly correlated with survival trends over time — the more dissimilar a species’ primary and logged forest niches are, the steeper the survival declines in logged forest. A recent paper from the same study system and study species found that the elevational ranges of species (and potential upslope movement; Girish & Srinivasan, 2022) explained changes in survival rates of species in primary but not in logged forest (Srinivasan & Wilcove, 2021). In logged forest therefore, rather than elevational ranges and range shifts, mismatches in the abiotic niche between primary and logged forest might better explain trends in survival. Although a reduction in the apparent survival of a species is possible through emigration alone, it is unlikely that the trends we observe arise from the increased emigration of individuals from the logged forest (see Srinivasan & Wilcove, 2021).

Such a relationship can occur in two ways. Firstly, an altered abiotic environment can interact directly with species’ physiology, especially on stress response mechanisms. It is likely that birds with more dissimilar temperature-humidity niche spaces (between primary and logged forest) face elevated levels of stress and, therefore, chronically higher levels of stress hormones over time (Wingfield et al., 2017). Heat-shock proteins (HSPs), which indicate long-term thermal stress, can be used to study this effect (Maak et al., 2003; Pusch et al., 2018). HSPs are also well suited for a mark-recapture methodology because other stress indicator hormones are shorter-lived and capture itself can lead to biased readings arising from temporary capture-related stress. Apart from HSPs, glucocorticoids (corticosterone) are hormones produced in response to stress in birds and can induce a wide range of metabolic cascades, such as the utilisation of stored nutrients for usage in physiological functions, changes in immunity and the prevention of oxidative damage (Marasco et al., 2017). A meta-analysis of avian response to logging revealed that species also have lowered immunity in logged forests in comparison with old-growth forests (Messina et al., 2018), and were likely to carry higher disease loads. Furthermore, glucocorticoid levels have been shown to have carry-over effects on population dynamics in many bird species (W. K. Hansen et al., 2016; Harms et al., 2015; Messina et al., 2020).

A change in abiotic conditions can also have indirect impacts on bird species; for instance, changes in temperature and humidity can impact the abundances and diversity of arthropod prey, which we demonstrate here. While one particular study of altered prey abundances of understory insectivores yielded little support for the prey-limitation hypothesis (Sekercioglu, 2011), the same species of birds often forage on arthropods at different trophic levels in primary versus logged forest (Hamer et al. 2015). The method of prey capture has been shown to be a consistent predictor of sensitivity to disturbance; for instance, sallying insectivores —those that capture flying prey while themselves in flight— were more sensitive to logging in African forests (Arcilla et al., 2015).

Our results suggest that the degree of niche dissimilarity is a strong predictor of survival declines of Eastern Himalayan birds in degraded habitats facing climate change. By using temperature and humidity loggers, this study demonstrates the applicability of niche estimation and comparison to reliably estimate the vulnerability of species to anthropogenic change. The IUCN Red List, widely used to set species-level conservation priorities, has been criticised for not accounting for climate-related threats to species (Thuiller et al., 2005). More generally, formal assessments of slow-acting threats such as climate change is a necessary but challenging step to kickstarting conservation action. Most often, however, by the time existing criteria such as population or range-size declines are observable at population levels, it may well be too late for effective conservation action (Hannah, 2012).

For effective conservation, it is essential that vulnerable species are identified as early as possible to maximise the chances of conservation success (Stanton et al., 2015). To this end, Climatic Niche Factor Analysis (CNFA) has been suggested as a tool to predict the differential vulnerability of various species to extinction (Rinnan & Lawler, 2019). However, the real-world applicability of niche modelling approaches to assess risk, especially in terms of demographic rates, is hitherto unexamined.

Our methodology of estimating temperature-humidity niche shifts that correlate with long-term survival declines in Eastern Himalayan bird species is one that potentially has widespread use in conservation management and planning. By providing managers and scientists with an inexpensive quantification method of the relative vulnerabilities of various species found in bird communities, such CNFA methodologies would enable targeted, prompt and judicious use of limited conservation resources for the species which are at the greatest risk of extinction due to global change drivers. Further research is needed to understand the physiological impacts of climate change and forest degradation/loss on tropical bird species. The integration of dietary and climatic requirements of species would also yield a more holistic understanding of the grave threat of anthropogenic change on tropical biodiversity worldwide.

## Acknowledgements

We thank S. Rai, B. Tamang, D.K. Pradhan, and M. Rai for their enthusiasm and professionalism in the field. We are grateful to the Arunachal Pradesh Forest Department for providing permission to conduct this study, and to Mr. Millo Tasser (IFS), Divisional Forest Officer of Shergaon Forest Division, for his continued support of our work. I. Glow, N. Tsering Monpa, and A. Pradhan helped greatly with logistics. Funding for this study came from the National Centre for Biological Sciences, International Foundation for Science, Princeton University, Princeton University’s Global Collaborative Network on Himalayan Biodiversity, and the High Meadows Foundation.

## Statement of authorship

AB and UM designed the study. UM collected the mark-recapture dataset from 2011 to 2021 and the temperature-humidity dataset in 2021. RC collected the arthropod community data. AB performed the survival, temperature-humidity and arthropod community analysis. AB wrote the first draft of the manuscript, and UM aided in its revision. UM supervised this work.

## Data accessibility statement

If accepted, all data supporting the results will be archived in an appropriate public repository and the data DOI will be included at the end of the article.

## Supplementary Material

**Figure S1:**
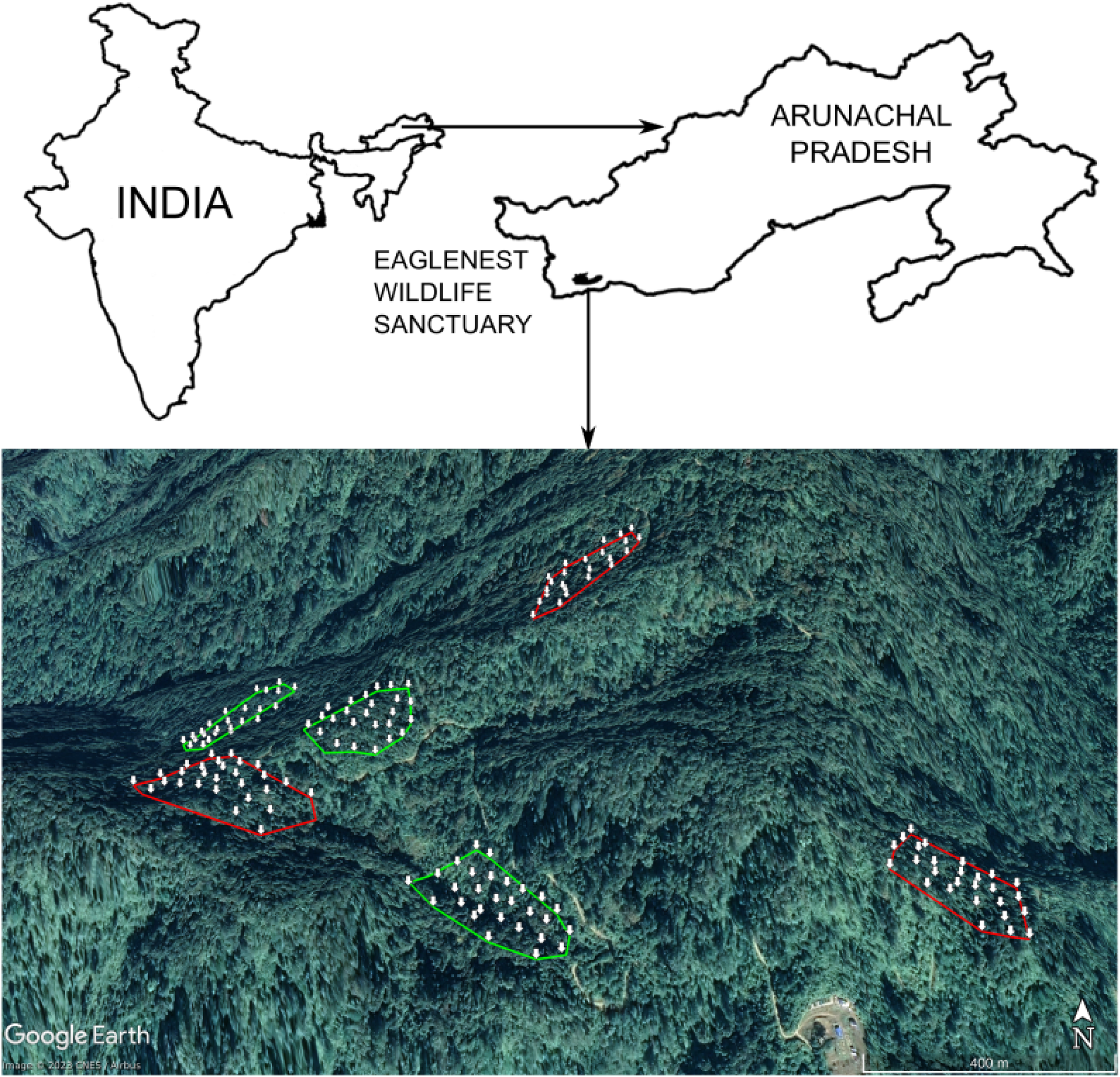
**A map of the study area in Eaglenest Wildlife Sanctuary, Arunachal Pradesh, India. Sampling plots in primary forest are outlined in green and plots in logged forest are outlined in red. White arrows show the locations of mist nets along with their corresponding temperature-humidity loggers.**

**Table S1:** **The common names and scientific names (Billerman et al., 2022) of the species selected for our analysis. All chosen species are breeders at our study site and had sufficient data for the modelling of survival trends and their temperature-humidity niches.**

| Common name | Scientific name |
| --- | --- |
| Black-throated Parrotbill | *Suthora nipalensis* |
| Rufous-capped babbler | *Cyanoderma ruficeps* |
| Yellow-throated Fulvetta | *Schoeniparus cinerea* |
| Rufous-winged Fulvetta | *Schoeniparus castaneiceps* |
| Golden-breasted Fulvetta | *Lioparus chrysotis* |
| Snowy-browed Flycatcher | *Ficedula hyperythra* |
| White-gorgetted Flycatcher | *Anthipes monileger* |
| Grey-cheeked Warbler | *Phylloscopus poliogenys* |
| Rusty-fronted Barwing | *Actinodura egertoni* |
| Golden Babbler | *Cyanoderma chrysaeum* |
| White-spectacled Warbler | *Phylloscopus intermedius* |
| Blyth's Leaf Warbler | *Phylloscopus reguloides* |
| Chestnut-crowned Laughingthrush | *Trochalopteron erythrocephalum* |
| Black-faced Warbler | *Abroscopus schisticeps* |
| Large Niltava | *Niltava grandis* |
| White-tailed Robin | *Myiomela leucura* |
| White-browed Piculet | *Sasia ochracea* |

**Table S2:** **Model parameters for the estimate and 95% confidence intervals around the estimate for the generalised linear model to study the relationship between body mass, time, habitat and niche overlap.**

| Model: | Z-score ~ Year * Habitat * Niche_Overlap |  |  |
| --- | --- | --- | --- |
| Parameter | Estimate | Lower CI bound | Upper CI bound |
| (Intercept) | 0.34 | 0.01 | 0.66 |
| Year | -0.04 | -0.08 | 0.01 |
| Niche_Overlap | -0.33 | -0.68 | 0.03 |
| Habitat_primary | -0.53 | -0.98 | -0.09 |
| Year:Niche_Overlap | 0.03 | -0.01 | 0.08 |
| Year:Habitat_primary | 0.04 | -0.02 | 0.1 |
| Niche_overlap:Habitat_primary | 0.52 | 0.04 | 1 |
| Year:Niche_Overlap:Habitat_primary | -0.04 | -0.11 | 0.03 |

**Table S3:** **Model parameters for the estimate and 95% confidence intervals around the estimate for the generalised linear model to study the relationship between body mass, time, habitat and niche overlap.**

| Model: | Phi(~Year * Habitat * Niche_Overlap)<br>p(~1) |  |  |
| --- | --- | --- | --- |
| Parameter | Estimate | Lower CI bound | Upper CI bound |
| Phi:(Intercept) | 0.57 | -0.46 | 1.59 |
| Phi:Year | -0.19 | -0.41 | 0.02 |
| Phi:Niche_Overlap | -0.87 | -1.99 | 0.24 |
| Phi:Habitat_primary | -1.98 | -3.41 | -0.55 |
| Phi:Year:Niche_Overlap | 0.24 | 0 | 0.47 |
| Phi:Year:Habitat_primary | 0.36 | 0.05 | 0.67 |
| Phi:Niche_Overlap:Habitat_primary | 2.27 | 0.71 | 3.83 |
| Phi:Year:Niche_Overlap:Habitat_primary | -0.37 | -0.71 | -0.04 |
| p:(Intercept) | -0.47 | -0.61 | -0.33 |

